# Holostean genomes reveal evolutionary novelty in the vertebrate immunoproteasome that have implications for MHCI function

**DOI:** 10.1101/2024.11.01.621518

**Authors:** Andi V. Barker, Kara B. Carlson, Dustin J. Wcisel, Ian Birchler De Allende, Ingo Braasch, Michael Fisk, Alex Dornburg, Jeffrey A. Yoder

## Abstract

Holosteans (gars and bowfins) have emerged as valuable models for understanding early vertebrate evolution, offering insights into diverse topics ranging from genomic architecture to molecular processes. These lineages also exhibit unusual features in their immune response, combining molecular elements seen in both tetrapods and ray-finned fishes. However, the immune repertoire of holosteans remains relatively unexplored. Here, we investigate the evolution of PSMB8, a core component of the immunoproteasome responsible for cleaving intracellular proteins into peptides for presentation by MHC class I molecules. We identify two holostean PSMB8 types—S type and K type—that are unique among vertebrates. These types likely cause significant biochemical changes to the S1 binding pocket involved in antigen cleavage which could result in the presentation of novel peptides by MHC class I. Integrating comparative analyses across major ray-finned fish lineages demonstrates that bowfins and gars independently evolved the PSMB8 S type within separate PSMB8 paralog lineages, while the PSMB8-K type is an evolutionary novelty found only in gars. Our results provide new perspectives into PSMB8 haplotypes and their role in peptide antigen processing, offering unique insights into the molecular evolution of the vertebrate immunity and antigen presentation.

## Introduction

Over the past decade, advances in genome sequencing have catalyzed a renaissance of interest in fish species originally dubbed “living fossils” by Darwin (Darwin 1861). Insights derived from the genomes of holosteans (gars and bowfins) have been particularly exciting, providing new perspectives into the molecular basis of early vertebrate diversification that include resolving key questions surrounding the evolution of genomic architecture (Braasch et al. 2016), developmental processes (Thompson et al. 2021), mobile elements (Braasch et al. 2016; Chalopin and Volff 2017; Mallik et al. 2023), and gene regulatory networks (Braasch et al. 2014; Braasch et al. 2016; Thompson et al. 2021) to name but a few. These findings are enabled by the phylogenetic position of holosteans as the sister lineage to the over 30,000 species of teleost fishes (Dornburg and Near 2021; Thompson et al. 2021), coupled with the observations that holosteans exhibit some of the slowest rates of molecular evolution of any vertebrate (Takezaki 2018; Brownstein et al. 2024)(Braasch et al. 2016)(Takezaki 2018; Brownstein et al. 2024). The demonstrated utility of holosteans for comparative studies has led to an emerging consensus that these lineages offer a genomic bridge between model teleost species and humans (Braasch et al. 2016; Dornburg and Near 2021). Although holosteans have provided groundbreaking insights into several long standing questions in vertebrate molecular evolution, molecular investigations into their immune repertoire have also revealed highly unusual features. Holosteans have been shown to possess molecular elements that appear as an amalgamation of features present in both tetrapods and ray-finned fishes. For example, holosteans possess an organization of the major histocompatibility complex (MHC) that is more similar to that in humans than teleosts such as zebrafish (Thompson et al. 2021). At the same time, holosteans also possess a diversity of immune functioning genes that are ray-finned fish specific, including putative natural killer receptor genes (Wcisel et al. 2017; Dornburg et al. 2021). This evolutionary unique combination of molecular elements now raises the question of how holosteans process antigens for immune surveillance.

For the adaptive immune response to function, a cell must be able to present “self” and “non-self” antigens on their surface for surveillance by immune cells. MHC genes play pivotal roles in this function, including the processing of peptide antigens and their subsequent extracellular presentation to other immune cells. This presentation is mediated by the inducible immunoproteasome, a multi-protein complex with proteolytic activities that cleave antigenic proteins into peptides for loading onto MHC class I for display to CD8^+^ T cells (Basler et al. 2013). However, the physical distance between proteasome genes and MHC class I or class II clusters comes with an evolutionary consequence. The genes responsible for cleaving peptides, processing peptides, and loading them on MHC class I are linked to MHC class I genes (in most non-mammals) or MHC class II genes (in most mammals) within the MHC locus (Van Kaer 2008; Ohta and Flajnik 2015; Palomar et al. 2021). Proteasome subunit genes in mammals are located far from the MHC class I genes, leading to less polymorphism in antigen processing genes and more diversity focused on class I haplotypes (McConnell et al. 2016). In contrast, non-mammalian species exhibit a highly conserved linkage between class I and proteasome genes, facilitating coevolution and coinheritance of alleles with compatible specificities for peptide cleavage, loading and presentation (Ohta et al. 2003; Kaufman 2013; McConnell et al. 2016; Veríssimo et al. 2023). Holostean genomes protease subunits are found near both MHC class I and class II genes (Thompson et al. 2021) suggesting a potential for evolutionary innovations in antigen processing genes that may be unique in the vertebrate Tree of Life.

Of the genes that constitute the immunoproteasome, *PSMB8* (LMP7/RING10/β_5i_) displays an unusual haplotypic feature of dimorphic alleles that reflect trans-species polymorphism, genetic variations that were present prior to speciation resulting in shared alleles in related species (Klein 1987; Nonaka et al. 2000; Klein et al. 2007; Azevedo et al. 2015). *PSMB8*’s well-documented trans-species polymorphism has made it an ideal candidate for investigating the evolutionary diversification of immunoproteasome components. Phylogenetic analyses have revealed PSMB8 types generally form two polymorphic clades with “A types” forming one lineage and “F types” forming a second lineage (referred to as “A” and “F” lineages, respectively), whose common ancestry coincides with the origin of vertebrate adaptive immunity (Tsukamoto et al. 2012). These polymorphisms are based on the chymotrypsin-like cleavage specificity that is determined by the 31st position residue in the S1 pocket of the mature protein (Unno et al. 2002; Tsukamoto et al. 2012) where substitutions are expected to affect the biochemical characteristics of the inner space, resulting in different enzymatic activity (Unno et al. 2002; Miura et al. 2010; Huang et al. 2013). The PSMB8A type encodes alanine or valine (A31 or V31) with small linear side chains and the PSMB8F type encodes phenylalanine or tyrosine (F31 or Y31) with larger aromatic side chains (Huang et al. 2013; Noro and Nonaka 2014). However, these A and F lineages have not been maintained across all vertebrates.

Trans-species polymorphism in PSMB8 is closely associated with the gene’s proximity to MHCI, leading to a reduction in allelic diversity in most tetrapods, likely due to the increased genomic distance between this gene and the MHCI cluster. In contrast, trans-species polymorphisms have been documented in lineages as diverse as sharks, cyprinids, salmonids, bichirs, and bony tongues (Miura et al. 2010; Fujito and Nonaka 2012; Tsukamoto et al. 2012; McConnell et al. 2016; Grimholt and Lukacs 2021). However, clade-specific losses have resulted in major groups of organisms losing either the A or F lineage, and then some of these lineages independently gaining lost function through mutation at the 31st position. For example, both tetrapods and acanthomorphs have lost the PSMB8F lineage. In a striking case of convergence, both *Oryzias* ricefish and *Xenopus* frog species have independently regained the PSMB8A “type” via mutation of the codon for the 31st position (Nonaka et al. 2000; Miura et al. 2010). Similarly, Anguilliformes (eels and tarpons) have lost the A lineage, and some eels such as *Conger* have independently regained A “type” function through a mutation in the F lineage (Noro and Nonaka 2014). Given the potential for clade specific losses and reversals of function, understanding the evolutionary history of holostean PSMB8 and identifying any putative novel genotypes necessitates an investigation of not just of holostean lineages, but also all other major ray-finned fish lineages to appropriately place PSMB8 lineages into their phylogenetic context.

Here we conducted the first investigation into the holostean immunoproteasome. Our analyses revealed two novel PSMB8 types in holosteans that are unique in the vertebrate Tree of Life, with residues in the S1 binding pocket that likely result in significant biochemical alterations and thus the properties of peptides displayed by MHC Class I. To place these types into a comparative framework, we integrated holostean PSMB8 sequences with an extensive survey of PSMB8 trans-species polymorphisms across all major ray-finned fish lineages and RNAseq data from numerous gar and bowfin individuals. Our results revealed that gar and bowfin each have their own novel type that corresponds to asymmetric losses of holostean PSMB8 lineages and that these types are broadly occuring in wild populations. Collectively, our findings suggest that the unique PSMB8 types in holosteans contribute to a molecular basis of immune surveillance that is unlike that found in any other group of vertebrates, providing new insights into the evolutionary processes that shape antigen processing mechanisms.

## Results and Discussion

### Holostean PSMB8 types suggest novel biochemical functions

Across all vertebrates, only two PSMB8A types (A31 or V31) and two PSMB8F types (F31 or Y31) have been reported (Huang et al. 2013; Noro and Nonaka 2014). However, our investigation into the available genomes of eyetail bowfin, spotted gar, longnose gar and alligator gar (Braasch et al. 2016; Bi et al. 2021; Thompson et al. 2021; Mallik et al. 2023) as well as new transcriptomes from 10 ruddy bowfin, 6 eyetail bowfin, and 29 longnose gar (**Supplementary Figures S1-S2, Supplementary Tables S1-S2**) revealed residues in position 31 that do not correspond with the A or F type. A comparison of holostean PSMB8 sequences with other actinopterygians (**Supplementary Tables S3-S4**) reveal that all bowfin PSMB8 sequences fall within the F lineage, but individual haplotypes encode either a phenylalanine (F), or a serine (S) at the 31st position (**Figure 1 and Supplementary Figures S1 and S3**). Unlike F, S does not include an aromatic side chain, but rather possesses a smaller, linear hydroxyl side group. As this sequence is structurally different from the A type and F type forms of PSMB8, we define this sequence as PSMB8S or S type (**Supplementary Figure S4**). The presence of the S type in both extant bowfin species suggests that this type may have arisen in extinct halecomorph lineages prior to the most recent common ancestor of living bowfins. In contrast, longnose gars do not possess either F or a S at the 31st position. All gar sequences fall within the A lineage and individual gar haplotypes possess either an Alanine (A), Threonine (T) or Lysine (K) at position 31 (**Figure 1** and **Supplementary Figures S2 and S3).** Threonine is structurally similar to serine with a smaller, linear hydroxyl side group, so we also refer to this sequence as S type (**Supplementary Figure S4**). Lysine (K) is distinctly different from all other residues reported for the 31st position of PSMB8 as it provides a positive charge — we refer to this sequence as PSMB8K or K type (**Supplementary Figure S4**). The presence of the S type in three gar species and the K type in two gar species suggests that these types may span all extant gar lineages.

**Figure 1.**
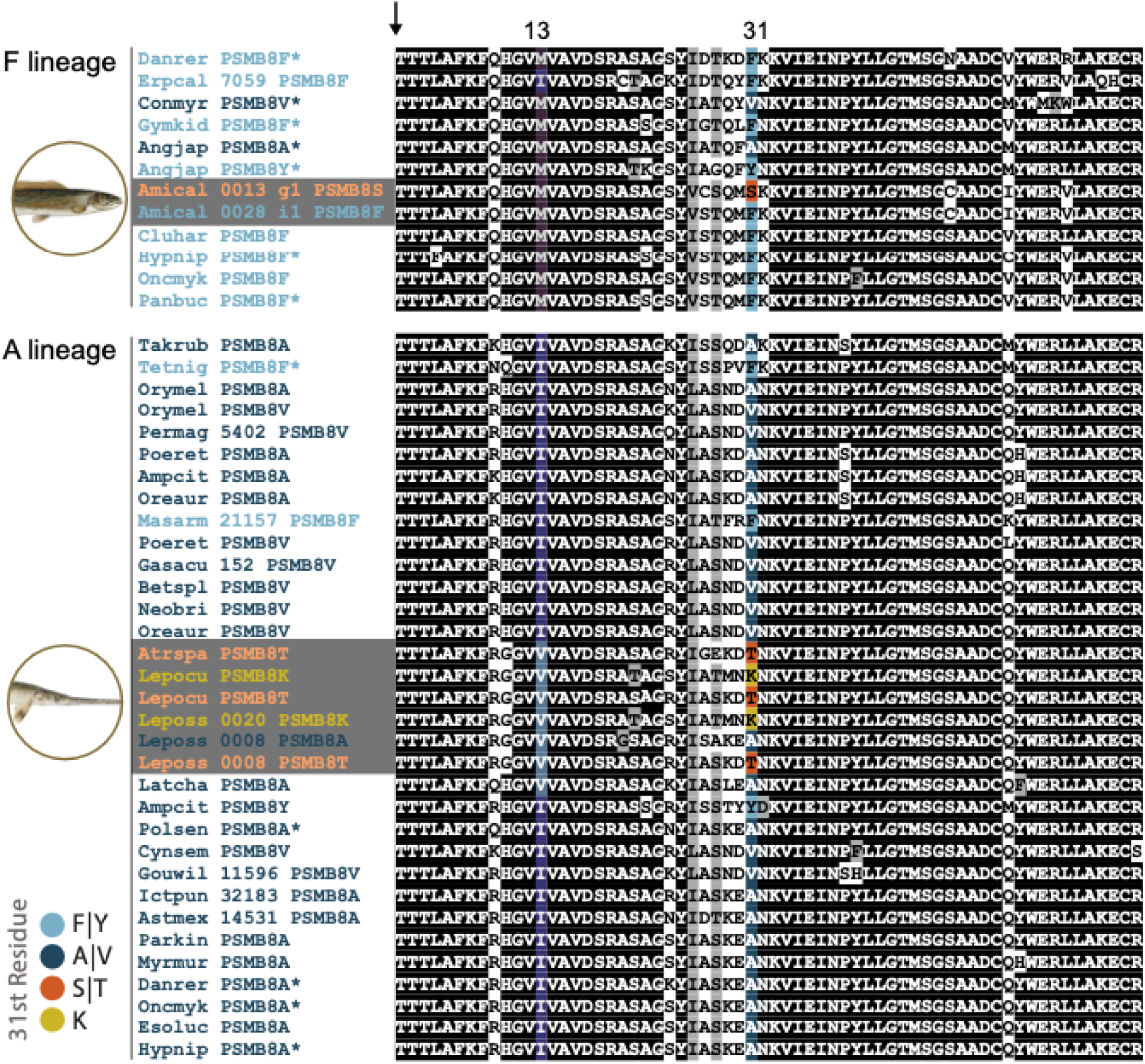
Alignment of the amino terminus of PSMB8 sequences. Representative A lineage and F lineage PSMB8 sequences from major Actinopterygian lineages. Sequences were aligned using Clustal Omega (Sievers and Higgins 2021). Positions that are ≥70% identical are shaded black and those that are structurally related are shaded gray. The predicted cleavage site which produces the mature protein is indicated with an arrow. The 13^th^ residue, in which M is described as predictive for the A lineage, and I as predictive for F lineage (Noro and Nonaka, 2014) are shaded violet or blue, respectively. Residues that do not match either lineage are additionally shaded zzz. Residue 31, which defines the PSMB8 type is shaded as defined in the key with taxon names shaded correspondingly. Sequence identifiers for ruddy bowfin (Amical) and longnose gar (Leposs) are indicated with gray shading. Sequences previously reported by Noro and Nonaka (2014) are indicated by asterisks (*). All sequences and display identifiers are provided in **Supplementary Tables S1-S4**. Alignments of full-length sequences are provided in **Supplementary Figure S3**.

To investigate how the S type and K type residues may impact PSMB8 function, we modeled all PSMB8 types from ruddy bowfin and longnose gar (**Figure 2**). The 31st residue of PSMB8 falls within the S1 pocket of the protein and mediates protease activity (Unno et al. 2002; Tsukamoto et al. 2012). As phenylalanine and alanine are structurally distinct, it has been proposed that the F type and A type possess recognize and cleave differing target proteins resulting in different peptides for display by MHC class I (Tsukamoto et al. 2012). However, it is unknown how the free hydroxyl group of the S type or the positively charged K type would alter the S1 binding pocket and cleavage properties. Our results demonstrate that these mutations are likely to have an effect on the binding pocket, with our space filling model indicating that S31 and T31 may occupy similar space in the binding pocket as A31 (**Figure 2**). As S and T can be phosphorylated by kinases, we additionally employed ScanSite Pro 4.0 (Obenauer et al. 2003) to determine if the 31st residue in PSMB8S or PSMB8T could be targets for phosphorylation (with the caveat that the database is built from mammalian kinases). The results of these scans for kinase target sequences suggests that T31 is not phosphorylated by any known kinase but that S31 could possibly be phosphorylated by Nek4 (albeit Nek4 only appeared in this scan using a low stringency screen). In humans, Nek4 is known to be involved in regulating cell cycle and DNA repair (Pavan et al. 2021). Whether this kinase has an alternate function or whether a kinase that has yet to be identified in holosteans phosphorylates S31 remains unknown.

**Figure 2.**
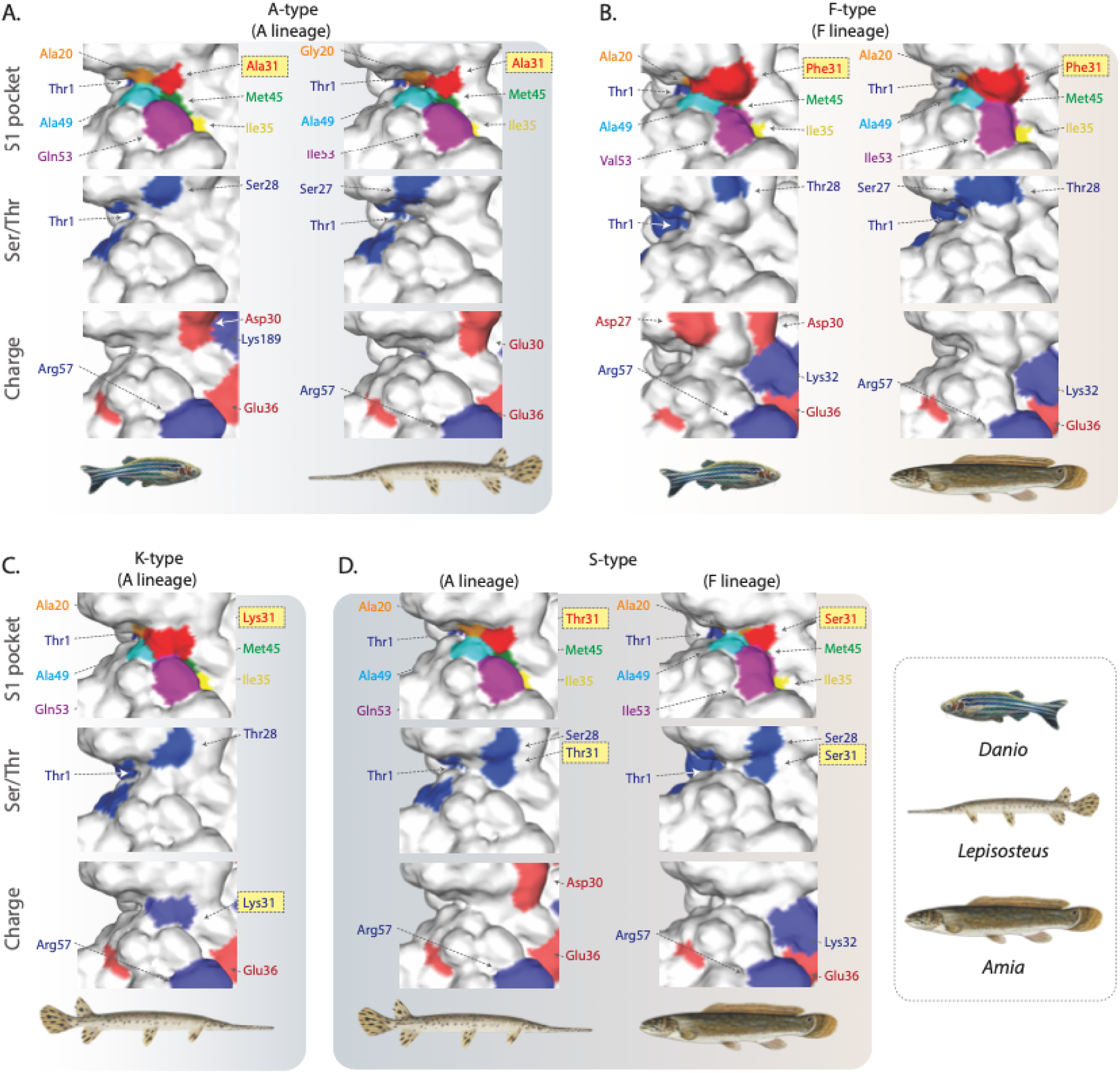
Predicted 3D structures of multiple PSMB8 types from zebrafish, ruddy bowfin and longnose gar. **(a)**. Structures of the PSMB8 S1 binding pocket were modeled **(A)** contrasting gar and zebrafish A types in the A lineage; **(B)** contrasting bowfin and zebrafish F types within the F lineage **(C)** evaluating the novel K type in longnose gar, and **(D)** contrasting the S types that occur in gar A lineage and bowfin F lineage. The same view of the binding pocket is shown with three different reference color sets. “S1 pocket” – the six residues forming the S1 pocket are color-coded as shown previously (Tsukamoto et al. 2012) with orange (20th position), red (31st), yellow (35th), green (45th), cyan (49th), and magenta (53rd), respectively and the catalytic threonine (Thr1) in blue. “Ser/Thr” – serines and threonines are shaded blue. “Charge” – positively charged residues are shaded blue and negatively charged residues are shaded red. Background shading differentiates between figure panels and fish images indicate representative taxa within each column.

Additionally, the PSMB8 K type is highly novel as it provides a positive charge within the S1 binding pocket. No other vertebrates are known to possess a charged residue in the 31st position of PSMB8. Our space filling model again indicates that the charged lysine would be centrally located within the binding pocket (**Figure 2**). The position of this charge may shift the recognition and cleavage preference of PSMB8 towards sequences with negatively charged residues. Future work to biochemically test this model offers an exciting research prospect with possible translational relevance.

Given that TAP1 (Transporter Associated with Antigen Processing 1) assists in transporting immunoproteasome-generated peptide fragments into the endoplasmic reticulum for loading onto MHC class I molecules (Shen and Rock 2006; Embgenbroich and Burgdorf 2018; Mantel et al. 2022), and that TAP1 is suggested to coevolve with *PSMB8* when they are linked (Ohta et al. 2003; Kaufman 2015), we investigated whether there are distinct forms of TAP1 associated with novel types of PSMB8 in gars and bowfins. With a focus on functional residues, we found no alternate forms of TAP1 specifically associated with these novel PSMB8 types. In fact the TAP1 functional residues were almost identical between holosteans and zebrafish, with only a conserved S to T substitution for residue 296 in longnose gar. This suggests that both holostean lineages utilize a mechanism similar to zebrafish for transporting peptides from the cytoplasm to the endoplasmic reticulum for loading onto MHCI (**Supplementary Figures S5-S6**).

### The Evolutionary History of Actinopterygian PSMB8

Eight diagnostic residues have been proposed for cataloging sequences into the PSMB8 lineages (A lineage/F lineage): residues in positions 13 (M/I), 99 (S/T), 147 (M/L), 150 (E/P), 156 (G/A), 188 (C/S), 189 (K/Q) and 194 (E/D) (Noro and Nonaka 2014). We determined the conservation of these residues (**Table 1** and **Supplementary Figure S3**), providing tentative support for a split between A and F types between gar and bowfin that correspond to overall alignment similarity. Seven of the eight residues from the F lineage are conserved in bowfin sequences, while five of the A lineage residues are conserved in gar sequences (**Table 1**). Although we did not observe 100% of these residues in gar and bowfin PSMB8 sequences matching the F or A lineages, our phylogenetic analysis (**Figure 3**) strongly supports that gars and bowfins have respectively maintained either the A or F lineage since their divergence over 200 million years ago (Near et al. 2012; Hughes et al. 2018).

**Figure 3.**
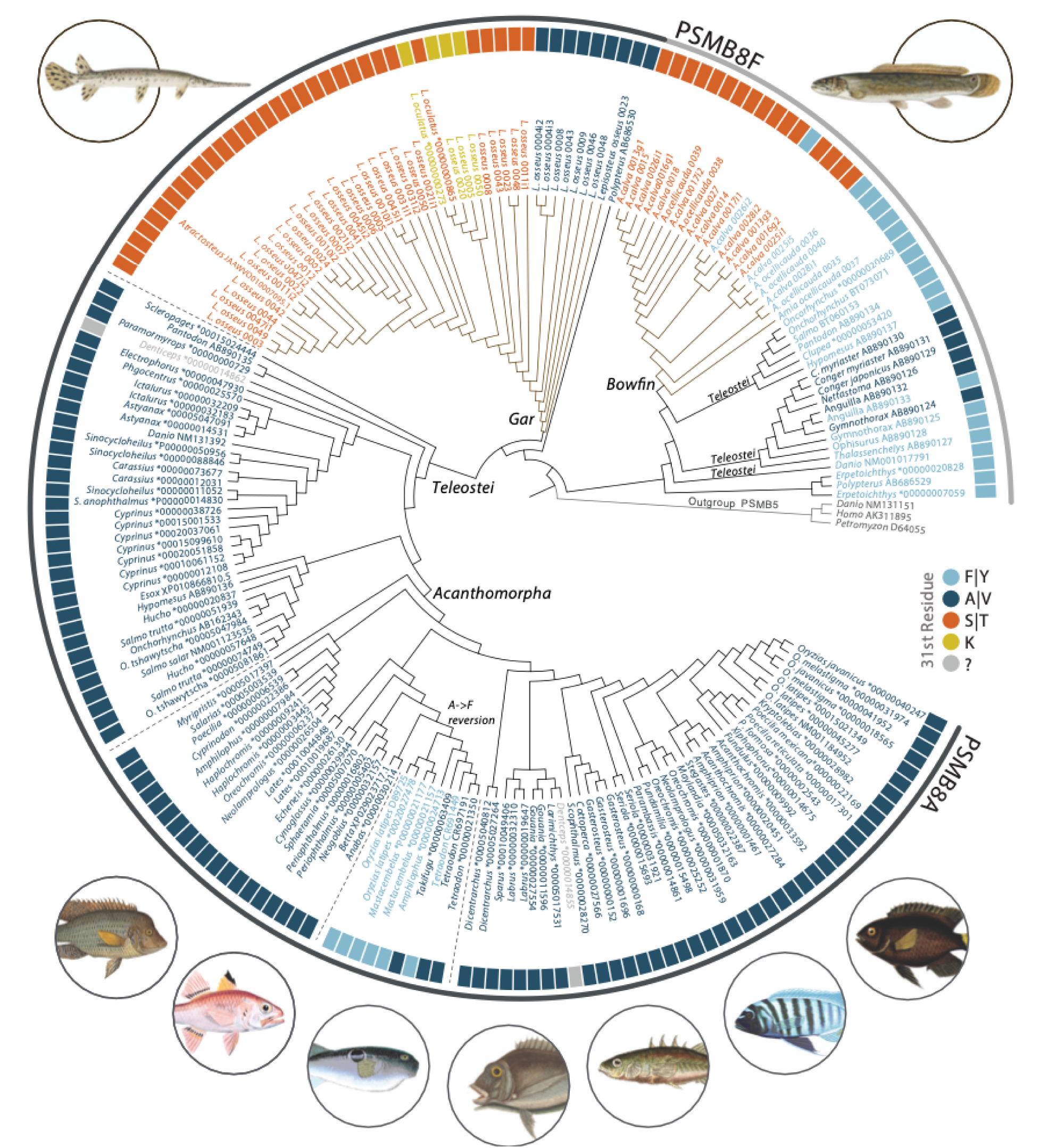
Phylogenetic relationships of PSMB8 sequences. Maximum likelihood tree topology of PSMB8 protein sequences across the evolutionary history of ray-finned fishes. Taxon names are color shaded based on the 31st position, with shadings corresponding to outer rectangles. Outerbands identify PSMB8F (gray) and PSMB8A (black) lineages. Holostean branches in the phylogeny are shaded to identify the most recent common ancestor of gar and bowfin PSMB8 sequences respectively. * in accession names indicate sequences from ENSEMBL. The functional equivalent of the 31st residue for denticle herring (*Denticeps*) PSMB8 is undefined (see main text and **Supplementary Figure S1**). PSMB5 sequences from human (*Homo*), zebrafish (*Danio*) and lamprey (*Petromyzon*) are included as an outgroup. Dashed lines correspond to three PSMB8A clades discussed in the text: Teleostei, Acanthomorpha, and the PSMB8A clade containing numerous independent acquisitions of the PSMB8F allele (A➜F reversion). Full length protein sequences are provided in **Supplementary Tables S1 – S4**. Fish images correspond to representative taxa within the phylogeny.

**Figure 4.**
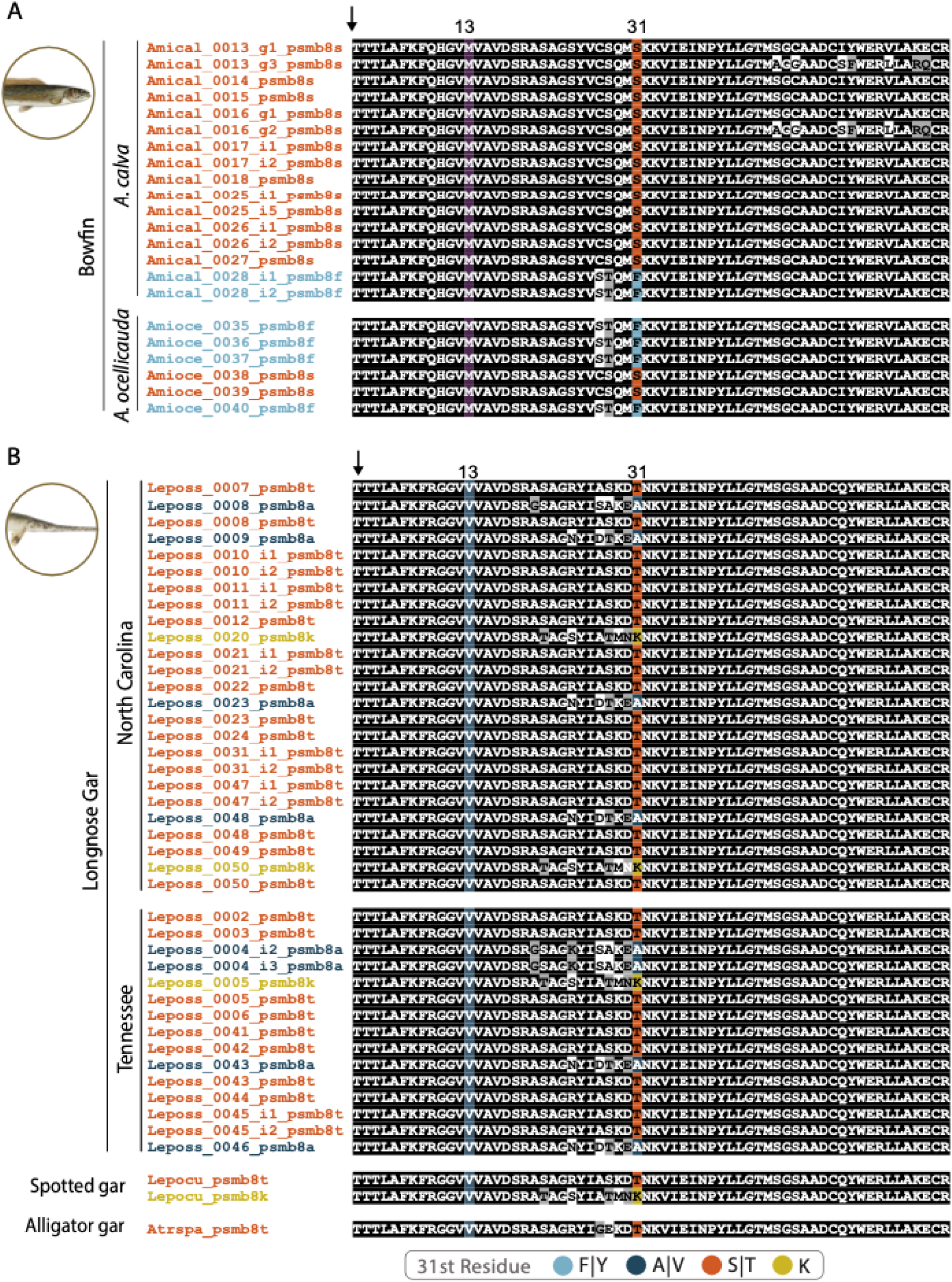
Alignment of the amino terminus of gar and bowfin PSMB8 sequences reveals variation at local spatial scales. Top: F lineage PSMB8 sequences from ruddy bowfin collected in North Carolina (*Amacal*) and from eyespot bowfin collected in Louisiana (*Amaoce*). Bottom, longnose gar (*Leposs*) sequences collected in either North Carolina or Tennessee, with additional gar sequences from spotted gar (*Lepocu*) and alligator gar (*Atrspa*) for comparison. Sequences were aligned using Clustal Omega (Sievers and Higgins 2021). Positions that are ≥70% identical are shaded black and those that are structurally related are shaded gray. The predicted cleavage site which produces the mature protein is indicated with an arrow. The 13^th^ residue, in which M is described as predictive for the A lineage (Noro and Nonaka, 2014), is shaded in violet. I is predictive for the F lineage (Noro and Nonaka, 2014), however no lineages possess an I. Residues that do not match either lineage are additionally shaded light blue. Residue 31, which defines the PSMB8 type is shaded as defined in the key with taxon names shaded correspondingly. Alignments of full-length sequences and a collection site details are provided in **Supplementary Figures S1, S2, S7 and Supplementary Tables S7-S8**.

**Table 1.** Diagnostic residues for PSMB8 lineages in gars and bowfin*.

| Position | A lineage | Gars | Bowfin | F lineage |
| --- | --- | --- | --- | --- |
| 13 | I | V | M | M |
| 99 | S | S | T | T |
| 147 | M | L | L | L |
| 150 | E | E | P | P |
| 156 | G | G | A | A |
| 188 | C | C | S | S |
| 189 | K | Q | Q | Q |
| 194 | E | E | E | D |

\* Diagnostic residues defined by Noro and Nonaka (2014).

To gain additional insights into the distribution of PSMB8 types in wild populations of holosteans, we placed our transcriptomic analyses of longnose gar, ruddy bowfin, and eyetail bowfin into the geographic context of collection sites from water bodies in the states of North Carolina (NC), Tennessee (TN), and Louisiana (LA) (**Supplementary Figures S1-S2, S7 and Supplementary Tables S7 – S8**). Phylogenetic analyses strongly support that all identified PSMB8 sequences from 10 ruddy bowfin and 6 eyetail bowfin fall within the F lineage (six F type sequences and 16 S type sequences; **Figure 3 and Supplementary Figure S1**). Similarly, we find that all identified PSMB8 sequences from 29 longnose gar fall within the A lineage (eight A type sequences, 24 S type (T31) sequences, and three K type sequences (**Figure 3** and **Supplementary Figure S2**). No other residues at position 31 were identified from these individuals, further suggesting that there has been no re-establishment of the A type in the bowfin or of the F type in longnose gar. Within our RNAseq data we identified A, T, and K types in longnose gar from both North Carolina and Tennessee. We additionally find F and S types in ruddy bowfin from North Carolina and eyetail bowfin from Louisiana. This result highlights that the S type in holosteans is independently derived in the two PSMB8 lineages, providing a striking example of potential functional convergence.

Considering the evolutionary history of the A and F lineages in the context of vertebrate evolution, there is a marked asymmetry in the maintenance of these lineages. The F lineage has been lost in most jawed vertebrates with the exception of cartilaginous fishes and non-acanthomorph ray-finned fishes. However, there are numerous examples of the F alleles being independently acquired within vertebrate A lineages. Our results increase the number of species within a clade of acanthomorphs that acquired F type alleles within the A lineage. Noro and Nonaka (2014) identified this independent acquisition in Atheriniformes (*Oryzias*) and Acanthuriformes (*Tetraodon* and *Takifugu*) and here, we report this independent gain in Synbranchiformes (*Mastacembelus*) and Blenniformes (*Amphilophus*) (**Figure 3**, **Table 2**). These results add new examples to documented cases of F alleles arising within the A lineage to cases documented in other vertebrates including amphibians (e.g., *Xenopus*), and reptiles (e.g., alligator, gecko, turtle) (Huang et al. 2013; Noro and Nonaka 2014). Our results are in line with the expectation that there is an asymmetry to the maintenance of the A lineage in vertebrates, and suggest that the F lineage was lost prior to the most recent common ancestor of acanthomorphs, a group that represents one out of every four living vertebrates (**Figure 3**). Acanthomorphs have been suggested to have experienced changes in the molecular organization of other immune related loci in the wake of major extinction events during the Cenozoic (Carlson et al. 2023), a hypothesis that is in line with growing recognition that extinction events can have a profound impact on genomic evolution (Berv et al. 2024). Our results suggest the F lineage may have followed a similar evolutionary trajectory. Further analyses of the immunoproteasome of all acanthomorphs are warranted to test this further.

**Table 2.**
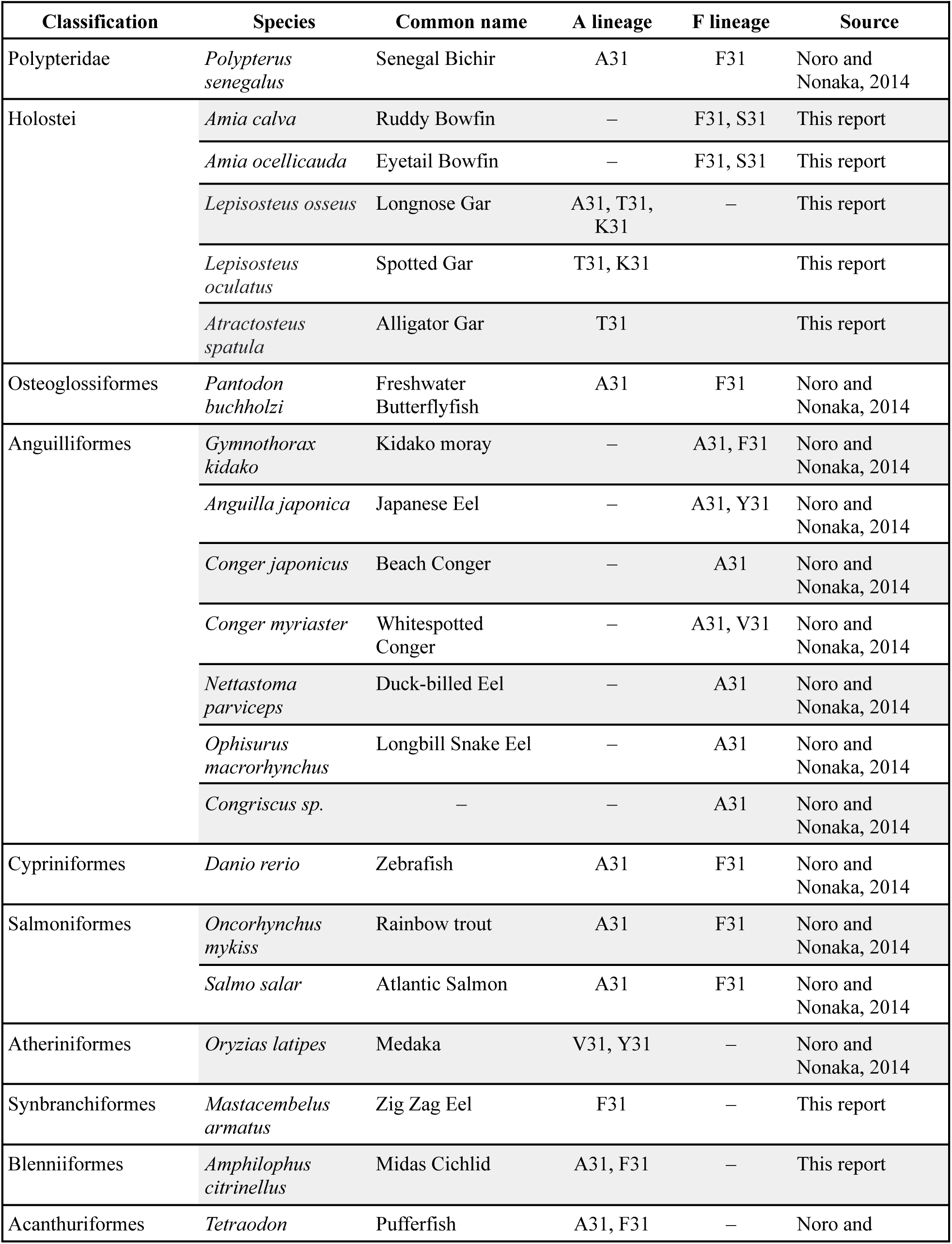
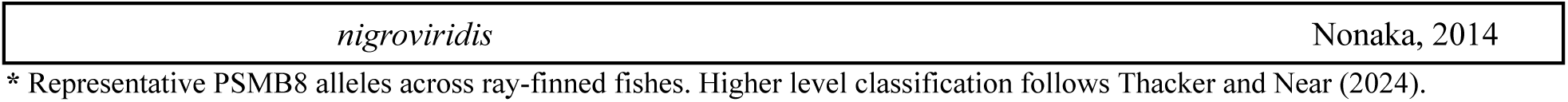
Distribution of PSMB8 alleles in ray-finned fishes*.

| Classification | Species | Common name | A lineage | F lineage | Source |
| --- | --- | --- | --- | --- | --- |
| Polypteridae | *Polypterus senegalus* | Senegal Bichir | A31 | F31 | Noro and Nonaka, 2014 |
| Holostei | *Amia calva* | Ruddy Bowfin | – | F31, S31 | This report |
|  | *Amia ocellicauda* | Eyetail Bowfin | – | F31, S31 | This report |
|  | *Lepisosteus osseus* | Longnose Gar | A31, T31, K31 | – | This report |
|  | *Lepisosteus oculatus* | Spotted Gar | T31, K31 |  | This report |
|  | *Atractosteus spatula* | Alligator Gar | T31 |  | This report |
| Osteoglossiformes | *Pantodon buchholzi* | Freshwater Butterflyfish | A31 | F31 | Noro and Nonaka, 2014 |
| Anguilliformes | *Gymnothorax kidako* | Kidako moray | – | A31, F31 | Noro and Nonaka, 2014 |
|  | *Anguilla japonica* | Japanese Eel | – | A31, Y31 | Noro and Nonaka, 2014 |
|  | *Conger japonicus* | Beach Conger | – | A31 | Noro and Nonaka, 2014 |
|  | *Conger myriaster* | Whitespotted Conger | – | A31, V31 | Noro and Nonaka, 2014 |
|  | *Nettastoma parviceps* | Duck-billed Eel | – | A31 | Noro and Nonaka, 2014 |
|  | *Ophisurus macrorhynchus* | Longbill Snake Eel | – | A31 | Noro and Nonaka, 2014 |
|  | *Congriscus sp.* | – | – | A31 | Noro and Nonaka, 2014 |
| Cypriniformes | *Danio rerio* | Zebrafish | A31 | F31 | Noro and Nonaka, 2014 |
| Salmoniformes | *Oncorhynchus mykiss* | Rainbow trout | A31 | F31 | Noro and Nonaka, 2014 |
|  | *Salmo salar* | Atlantic Salmon | A31 | F31 | Noro and Nonaka, 2014 |
| Atheriniformes | *Oryzias latipes* | Medaka | V31, Y31 | – | Noro and Nonaka, 2014 |
| Synbranchiformes | *Mastacembelus armatus* | Zig Zag Eel | F31 | – | This report |
| Blenniiformes | *Amphilophus citrinellus* | Midas Cichlid | A31, F31 | – | This report |
| Acanthuriformes | *Tetraodon* | Pufferfish | A31, F31 | – | Noro and |

\* Representative PSMB8 alleles across ray-finned fishes. Higher level classification follows Thacker and Near (2024).

Relative to the A lineage, the F lineage appears sequence-depauperate. However, our results also provide rare examples in which the F lineage is maintained and forms the substrate for evolutionary innovation. To date, only Anguilliformes were known to have lost the A lineage, with PCR-based amplification of single individuals suggesting they possess only the F type (e.g., *Congriscus sp*. [on NCBI as *Thalassenchelys*] and *Ophisurus macrorhynchus*), both the F and A types through mutations in the F lineage (e.g., *Gymnothorax kidako* and *Anguilla japonica*), or only the A type (e.g., *Conger myriaster*, *Conger japonicus*, and *Nettastoma parviceps* (Noro and Nonaka 2014). Our work reveals that the independent gain of an alternate PSMB8 type from the F lineage in Angulliforms is mirrored in bowfins that have lost the A lineage and derived the novel S type from the F lineage. Additionally, our analyses identified unusual features of two PSMB8 sequences from denticle herring (*Denticeps clupeoides*), within the A lineage (**Figure 3, Supplementary Figure S3**). These sequences may reflect paralogs, and we tentatively name them PSMB8F and PSMB8S based on their alignment with other PSMB8 sequences, and not based on the actual 31^st^ position due to lack of type resolution deriving from insertions and deletions (**Supplemental Figure S3**). Future studies will be needed to resolve the structure-function relationships for these PSMB8 forms. Regardless, holosteans offer a unique perspective on the observed imbalance in extant sequence diversity between A and F lineages that sets the stage for future investigations aimed at identifying evolutionary trends in the immunoproteasome across all vertebrates.

## Conclusion

Our investigation into holostean immunoproteasome reveals novel PSMB8 residues at the 31st position in bowfins and gars, revealing the presence of unique S and K type sequences. Bowfin sequences clustered within the F lineage exhibited a serine (S) residue in addition to an F type, while gar sequences within the A lineage displayed either threonine (T) or lysine (K) in addition to the expected A type. Structural modeling suggests that these residues likely alter the S1 binding pocket’s biochemical characteristics, impacting peptide cleavage and thus the peptides displayed by MHC class I. Our phylogenetic analysis supports that gars and bowfins have independently maintained the A and F lineages, respectively, with no evidence of re-establishment of the other lineage. This asymmetry aligns with broader vertebrate trends, where the F lineage is typically lost, but the F type is sometimes regained through mutations in the A lineage, as observed in some acanthomorph fish, amphibians, and reptiles. The unique PSMB8 types in holosteans suggest a general feature of A lineage persistence over deep time scales but also highlight rare instances where the F lineage is maintained and innovated upon. Moreover, the independent gain of the S type in both lineages marks a striking example of convergence in the immunoproteasome. The distinctiveness of holostean PSMB8 types prompts new questions about the evolutionary trajectory of PSMB8 and sets the stage for research on the functional implications of these unique PSMB8 types, especially their role in antigen presentation. Such research could have substantial implications for understanding the evolution of antigen display in the vertebrate immune system with translational relevance for human health.

## Methods

### Characterizing holostean PSMB8 sequence identity and structure

PSMB8 has been annotated in the spotted gar genome (Braasch et al. 2016). We verified this annotation by comparing sequence similarity to known actinopterygian PSMB8 sequences and used spotted gar PSMB8 as a query for BLAST searches in the genomes of eyetail bowfin (*Amia ocellicauda*, (Thompson et al. 2021)), longnose gar (Lepisosteus osseus, Mallik et al. 2023), and alligator gar (Atractosteus spatula, Bi et al. 2021). We then employed these longnose gar and eyetail bowfin reference sequences as queries for BLAST searches of our assembled transcriptomes (see below) from both species of bowfin and longnose gar, respectively. The retrieved sequences were aligned using Clustal Omega, and the 31st residue of the mature PSMB8 protein was determined for each sequence. To model the three-dimensional structures of these mature PSMB8 types, homology modeling was performed using the automated mode of the SWISS-MODEL server (Bordoli et al. 2009) via Expasy (Duvaud et al. 2021), following methods previously described by Tsukamoto et al. (2012).

Specifically, the structure of bovine PSMB5 (Protein Data Bank ID code: 1IRU) (Unno et al. 2002) was used as a template for modeling PSMB8 types, including ruddy bowfin S type (**Supplementary Table S1**: Amical 0013 g1 psmb8s) and F type (**Supplementary Table S1**: Amical 0028 i1 psmb8f), and longnose gar A type (**Supplementary Table S2**: Leposs 0008 psmb8a), S type (**Supplementary Table S2**: Leposs 0008 psmb8t), and K type (**Supplementary Table S2**: Leposs 0020 psmb8k). These types were compared to models of zebrafish A type (Genbank NP_571467.3) and F type (Genbank NP_001017791.1).

### Assessing sequence variation in TAP1

Using the human TAP1 sequence (GenBank ID: NP_000584.3) as a query, we identified putative TAP1 sequences in bowfin and longnose gar by searching against transcriptome assemblies on a custom internal BLAST server (Priyam et al. 2019). TAP1 sequences missing start and stop codons were excluded. In bowfin, this analysis revealed two forms of TAP1 in both species. The presence of these two TAP1 sequences in bowfin was further validated by BLAST analysis of the predicted cDNAs against the *Amia ocellicauda* reference genomes. Full length TAP1 cDNAs for longnose gar and bowfin were then translated via the ExPASy web server “translate” tool (Gasteiger et al. 2003) and aligned using Clustal Omega (Sievers and Higgins 2021). The essential residues required for TAP1 were determined based on the review by Lehnert and Tampé (2017). Regions essential for peptide sensing, binding, substrate specificity, and functionality were inspected and compared to assess if holosteans possess unique residues associated with their novel PSMB8 types (**Supplementary Figures S5-S6**).

### Transcriptomic analysis of wild holostean populations

All work with live animals was performed in accordance with relevant institutional and national guidelines and regulations, and was approved by the Institutional Animal Care and Use Committee of North Carolina State University (protocol 17-127-O) or Michigan State University (protocol 10/16-179-00). Bowfin and longnose gar were collected from waterways in North Carolina (NC), Tennessee (TN), and Louisiana (LA) (**Supplementary Figure S7**, **Supplementary Tables S7 and S8**). Immune tissues (gill, spleen, intestine) were dissected from 29 longnose gar, 10 ruddy bowfin and 6 eyetail bowfin (see below) and RNA was extracted from each individual tissue following manufacturer protocols (RNeasy kit; Qiagen, Germantown, MD). RNA was quantified using a NanoDrop 1000 (ThermoFisher, Waltham, MA) and Bioanalyzer (Agilent, Santa Clara, CA). In brief, mRNA was enriched using oligo(dT) beads, rRNA was removed using a Ribo-Zero kit (Epicentre, Madison, WI) and mRNA was randomly fragmented. Each RNA sample was diluted to 180 ng/µl and samples from the same individual were pooled for cDNA library preparation and sequencing which were performed by Novogen Corporation (Sacramento, CA). Next-gen sequencing (2 × 150 bp paired end reads) was performed on a NovaSeq 6000 instrument (Illumina, San Diego, CA). Adapter sequences and poor quality reads were filtered with Trimmomatic v34 (Bolger et al. 2014). Transcriptomes were de novo assembled using Trinity v2.11.0 (Grabherr et al. 2011). The confirmed spotted gar PSMB8 sequence (Ensembl ENSLOCP00000000273) was then used as a query for BLAST searches of the longnose gar and bowfin transcriptomes. Sequence hits with an e-value < –10 were retained for further analysis. PSMB8 protein sequences were inspected using Geneious Prime 2021.2 (https://www.geneious.com), and sequences lacking the consensus TTTL (Thr-Thr-Thr-Leu) sequence, which marks the start of the mature protein (Ferrington and Gregerson 2012; Noro and Nonaka 2014), or without an identifiable 31st amino acid were excluded from further analysis.

Bowfin collected in Louisiana were caught near the biogeographic break between the *Amia calva* and the newly delimited *Amia ocellicauda (Brownstein et al. 2022; Wright et al. 2022)*. Extensive genetic sampling along this drainage has revealed no occurrences of *Amia calva* (Brownstein et al. 2022), however as bowfin can tolerate saline conditions (Chipman 1959; Pearson and Ward 1972) it is possible an aberrant individual crossed a stretch of the Gulf of Mexico. As such we verified the identity of the species collected using COI barcode references for the two species (*Amia calva:* JN024760.1; *Amia ocellicauda:* KX145442.1), thereby ensuring correct identification.

### Placing Holostean PSMB8 Sequences into their Evolutionary Context

We augmented our sampling of PSMB8 sequences from holosteans with all annotated Actinopterygian sequences available in the Ensembl genome database (version 100). This sampling added our 51 Holostean sequences to 123 sequences from Acanthuriformes (n=8); Anguilliformes (n=11); Atheriniformes (n=17); Beryciformes (n=1); Blenniiformes (n=20); Carangiformes (n=7); Characiformes (n=3); Clupeiformes (n=3); Cypriniformes (n=16); Gobiiformes (n=3); Gymnotiformes (n=1); Labriformes (n=2); Osmeriformes (n=2); Osteoglossiformes (n=4); Perciformes (n=4); Polypteridae (n=4); Salmoniformes (n=12); Salmoniformes (n=12); Siluriformes (n=2); and Synbranchiformes (n=4). The above clade names follow the phylogenetic classification of ray-finned fishes (Dornburg and Near 2021; Near and Thacker 2024). Peptide sequences corresponding to intronic regions were removed from protein sequences for subsequent analyses. We utilized PSMB5 sequences from human, sea lamprey, and zebrafish to serve as outgroups (Hughes 1997; Bos 2005).

Full-length protein sequences were aligned using Clustal Omega (Sievers and Higgins 2021) and the 31^st^ residue of the mature PSMB8 protein was determined and used to define PSMB8 type (Schmidtke et al. 1996; Ferrington and Gregerson 2012; Noro and Nonaka 2014). Sequences which had an undefined residue at the 31^st^ position or were truncated and missing the 31^st^ residue were removed from the analyses. If multiple identical sequences were identified from the same species, only one was included in the analyses. In two cases, inspection of sequences from ENSEMBL revealed likely missannotation (e.g., *Esox lucius*, ENSELUP00000041989; *Betta splendens*, ENSBSLP00000026526). For these taxa, we conducted a search on NCBI genbank, finding alternative PSMB8 sequences (*Esox lucius*, XP_010866810.5; *Betta splendens,* XP_029023717.1) which were employed in all future analyses. Evolutionary relationships of PSMB8 sequences were inferred using maximum likelihood in IQ-TREE2 (Nguyen et al. 2015), with the best-fit model of AA substitution selected using ModelFinder (Kalyaanamoorthy et al. 2017). The candidate pool of substitution rates included all common amino acid exchange rate matrices (JTT, WAG, etc), protein mixture models such as empirical profile mixture models, as well as parameters to accommodate among-site rate variation (discrete gamma or free rate model). Node support was assessed via 1,000 ultrafast bootstrap replicates (Hoang et al. 2018). Phylogenetic analyses were used to determine the PSMB8 lineage of each sequence, with lineage designations defined in Noro and Nonaka (2014) serving as a reference.

## Acknowledgements

We thank Solomon David and Allyse Ferrara (Nicholls State University) for help with bowfin sample acquisition. We thank Martin Flajnik (University of Maryland) for helpful discussions about the immunoproteasome, and members of the Dornburg and Yoder labs for helpful comments during the early stages of this project. All fish images used in this manuscript were obtained from the public domain with the exception of the zebrafish image that is available for public use courtesy of the database for life science under a creative commons attribution 4.0 international license: doi.org/10.7875/togopic.2021.004.

## Contributions

DJW conceived the project; AVB, KBC and DJW datamined public databases; DJW, IB, MF, AD and JAY collected samples; KBC, DJW and IBDA completed transcriptome analyses; AVB, AD, KBC, DJW, and IBDA performed phylogenetic analyses; AD, AVB, KBC and JAY created graphics; AD, AVB, KBC, IBDA and JAY wrote the first draft of the manuscript; JAY and AD supervised the project. All authors read and approved the manuscript.

## Data availability

Raw reads and computationally assembled transcriptome sequences were deposited onto NCBI under the BioProject accession number PRJNA990750.

