## Supplemental Figures for "Holostean genomes reveal evolutionary novelty in the vertebrate immunoproteasome that have implications for MHCI function"

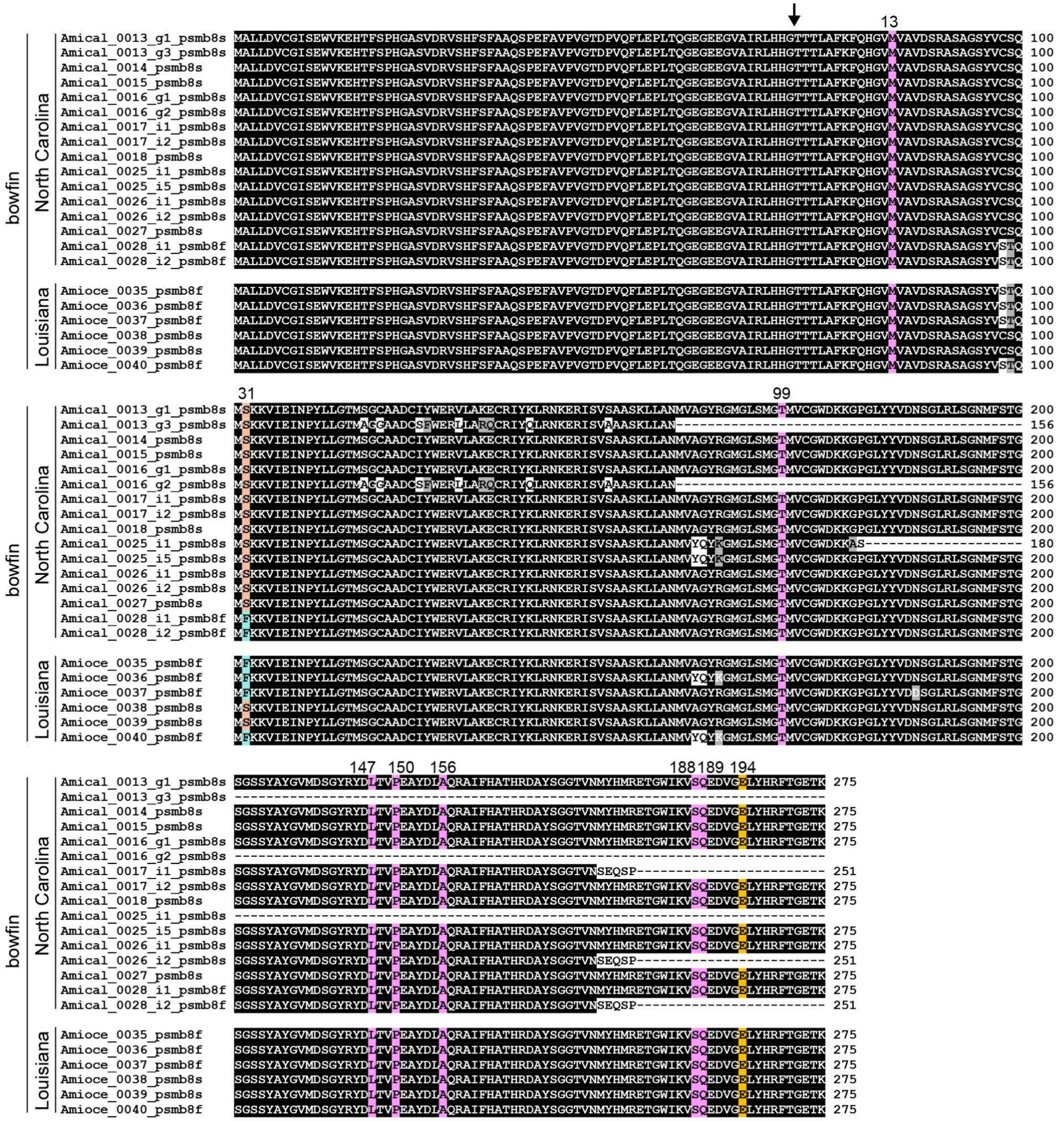

**Supplementary Figure S1. Bowfin PSMB8 sequences are F lineage.**

Full-length PSMB8 sequences from ruddy bowfin (North Carolina) and eyespot bowfin (Louisiana) were aligned using Clustal Omega (Sievers and Higgins, 2021). Positions that are ≥70% identical are shaded black and those that are structurally related are shaded gray. The predicted cleavage site which produces the mature protein is indicated with an arrow. The eight residues described as predictive for the A and F lineages (Noro and Nonaka 2014) are shaded gold and pink, respectively. Residues that do not match either lineage are shaded blue. Residue 31, which defines the PSMB8 type is shaded orange (S type) or aqua (F type). All sequences and display identifiers are provided in **Supplementary Table S1.**

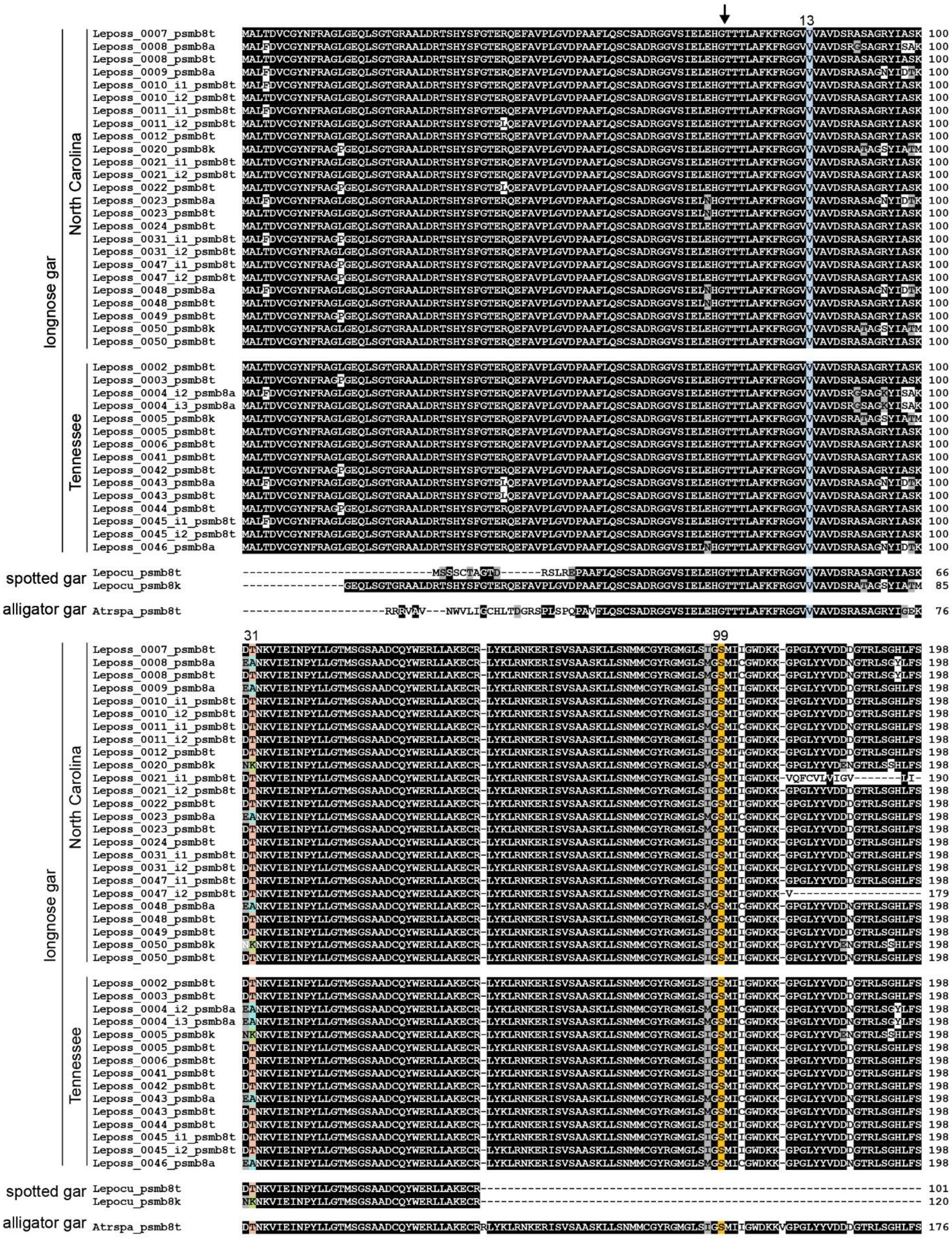
longnose gar
North Carolina
Tennessee
spotted gar
alligator gar
13
31
99

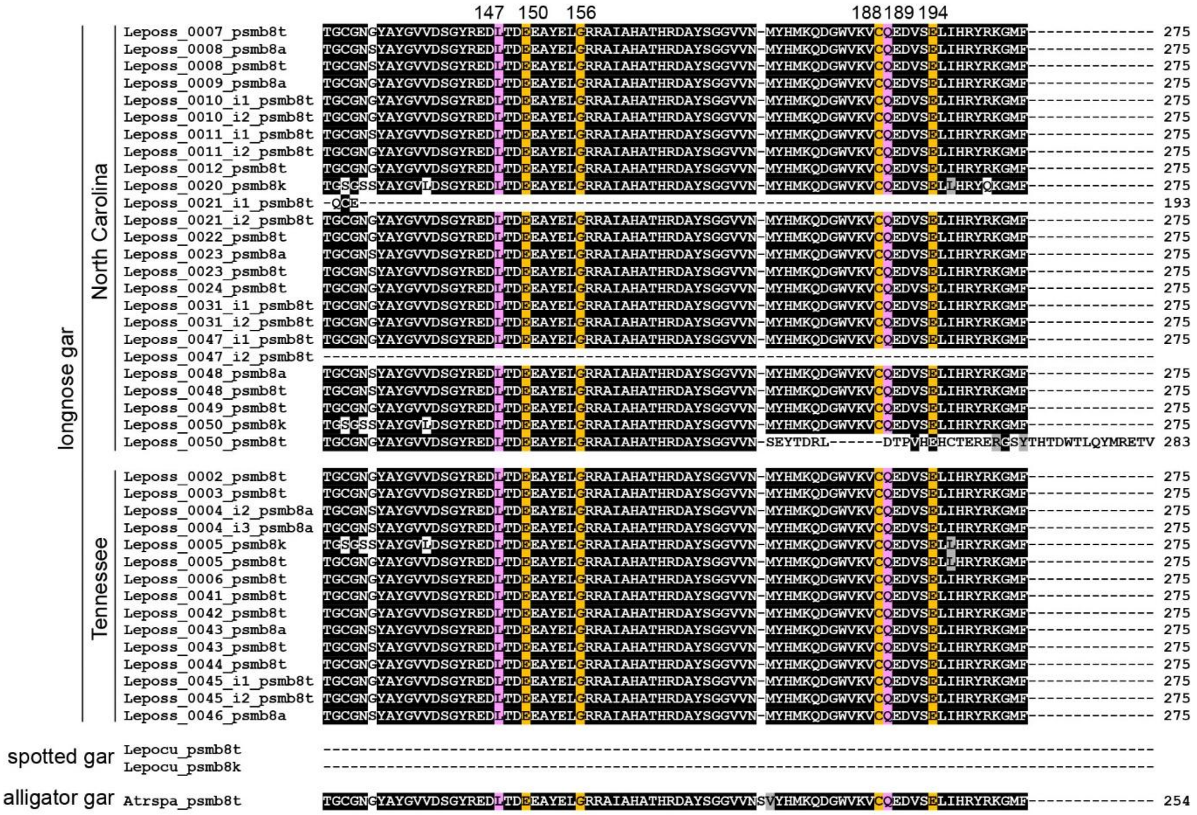

**Supplementary Figure S2. Gar PSMB8 sequences are A lineage.**

Full-length longnose gar PSMB8 sequences from North Carolina and Tennessee were aligned with reference sequences from spotted gar (Braasch et al. 2016) and alligator gar (Bi et al. 2021) using Clustal Omega (Sievers and Higgins 2021). Positions that are ≥70% identical are shaded black and those that are structurally related are shaded gray. The predicted cleavage site which produces the mature protein is indicated with an arrow. The eight residues described as predictive for the A and F lineages (Noro and Nonaka 2014) are shaded gold and pink, respectively. Residues that do not match either lineage are shaded blue. Residue 31, which defines the PSMB8 type is shaded orange (T type), aqua (A type) or green (K type). All sequences and display identifiers are provided in **Supplementary Table S2**.

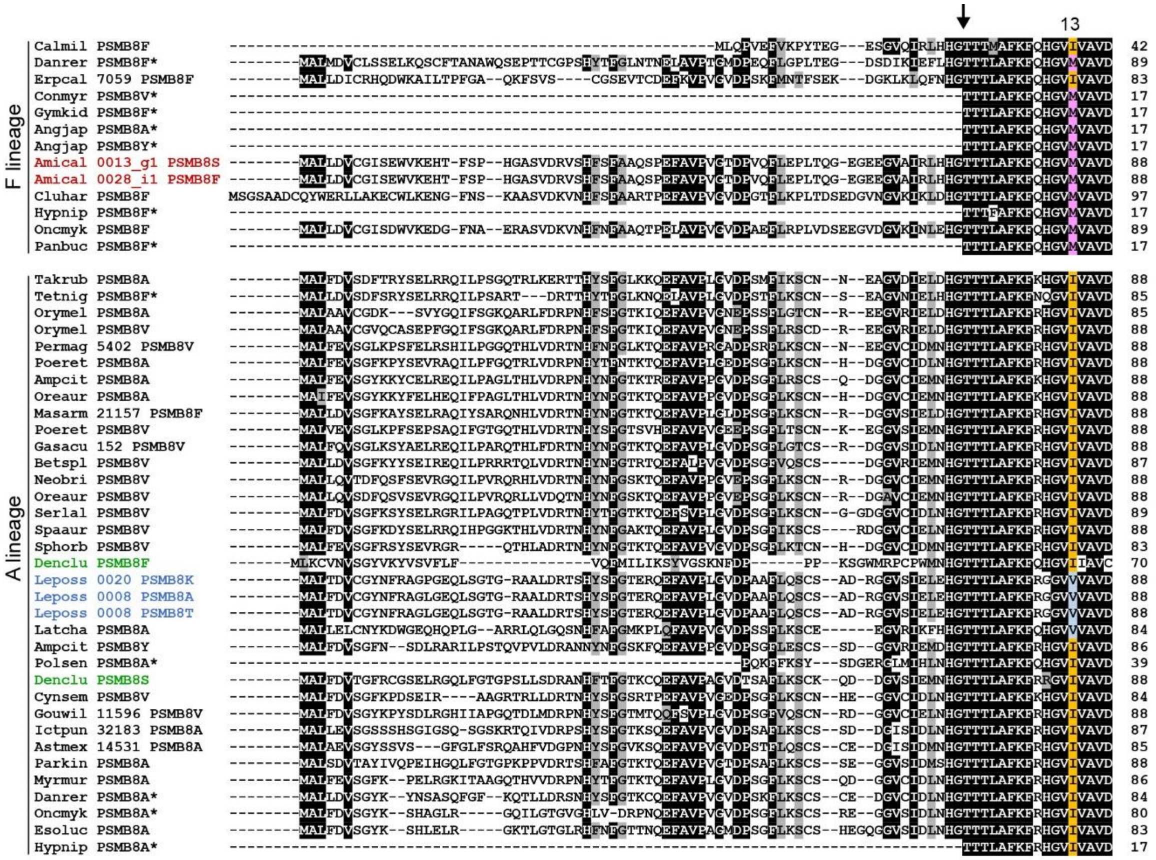
13
F lineage
Calmil PSMB8F
Danrer PSMB8F*
Erpcal 7059 PSMB8F
Conmyr PSMB8V*
Gymkid PSMB8F*
Angjap PSMB8A*
Angjap PSMB8Y*
Amical 0013_g1 PSMB8S
Amical 0028_i1 PSMB8F
Cluhar PSMB8F
Hypnip PSMB8F*
Oncmyk PSMB8F
Panbuc PSMB8F*
A lineage
Takrub PSMB8A
Tetnig PSMB8F*
Orymel PSMB8A
Orymel PSMB8V
Permag 5402 PSMB8V
Poeret PSMB8A
Ampcit PSMB8A
Oreaur PSMB8A
Masarm 21157 PSMB8F
Poeret PSMB8V
Gasacu 152 PSMB8V
Betspl PSMB8V
Neobri PSMB8V
Oreaur PSMB8V
Serlal PSMB8V
Spaaur PSMB8V
Sphorb PSMB8V
Denclu PSMB8F
Leposs 0020 PSMB8K
Leposs 0008 PSMB8A
Leposs 0008 PSMB8T
Latcha PSMB8A
Ampcit PSMB8Y
Polsen PSMB8A*
Denclu PSMB8S
Cynsem PSMB8V
Gouwil 11596 PSMB8V
Ictpun 32183 PSMB8A
Astmex 14531 PSMB8A
Parkin PSMB8A
Myrmur PSMB8A
Danrer PSMB8A*
Oncmyk PSMB8A*
Esoluc PSMB8A
Hypnip PSMB8A*

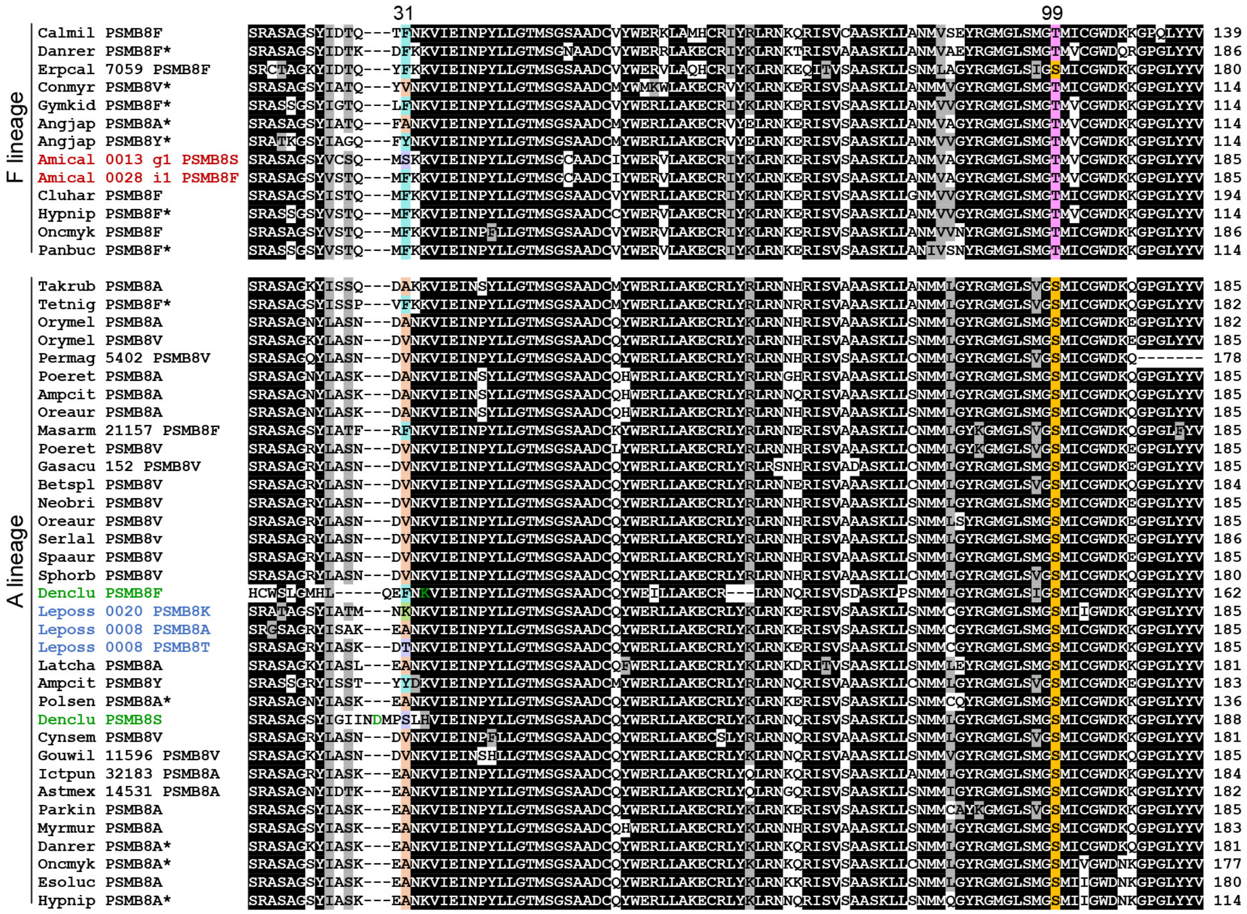
31
99
F lineage
A lineage
Calmil PSMB8F 139
Danrer PSMB8F* 186
Erpcal 7059 PSMB8F 180
Conmyr PSMB8V* 114
Gymkid PSMB8F* 114
Angjap PSMB8A* 114
Angjap PSMB8Y* 114
Amical 0013 g1 PSMB8S 185
Amical 0028 i1 PSMB8F 185
Cluhar PSMB8F 194
Hypnip PSMB8F* 114
Oncmyk PSMB8F 186
Panbuc PSMB8F* 114
Takrub PSMB8A 185
Tetnig PSMB8F* 182
Orymel PSMB8A 182
Orymel PSMB8V 185
Permag 5402 PSMB8V 178
Poeret PSMB8A 185
Ampcit PSMB8A 185
Oreaur PSMB8A 185
Masarm 21157 PSMB8F 185
Poeret PSMB8V 185
Gasacu 152 PSMB8V 185
Betspl PSMB8V 184
Neobri PSMB8V 185
Oreaur PSMB8V 185
Serlal PSMB8v 186
Spaaur PSMB8V 185
Sphorb PSMB8V 180
Denclu PSMB8F 162
Leposs 0020 PSMB8K 185
Leposs 0008 PSMB8A 185
Leposs 0008 PSMB8T 185
Latcha PSMB8A 181
Ampcit PSMB8Y 183
Polsen PSMB8A* 136
Denclu PSMB8S 188
Cynsem PSMB8V 181
Gouwil 11596 PSMB8V 185
Ictpun 32183 PSMB8A 184
Astmex 14531 PSMB8A 182
Parkin PSMB8A 185
Myrmur PSMB8A 183
Danrer PSMB8A* 181
Oncmyk PSMB8A* 177
Esoluc PSMB8A 180
Hypnip PSMB8A* 114

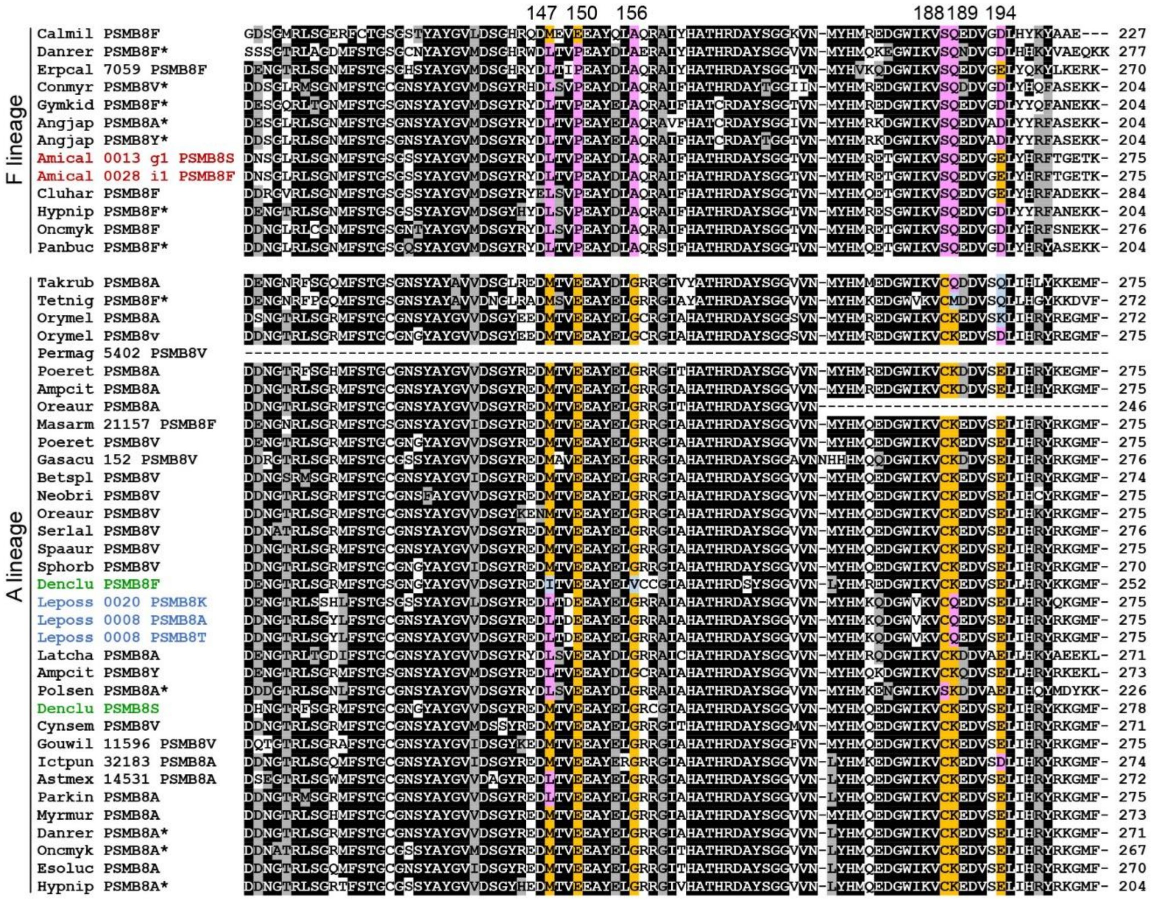

**Supplementary Figure S3. Alignment of select PSMB8 sequences.**

A lineage and F lineage PSMB8 sequences from major Actinopterygian lineages were aligned using Clustal Omega (Sievers and Higgins 2021). Efforts were made to include major radiations as well as examples of unusual sequence types (residue 31). Positions that are ≥70% identical are shaded black and those that are structurally related are shaded gray. The predicted cleavage site which produces the mature protein is indicated with an arrow. The eight residues described as predictive for the A and F lineages - 13, 99, 147, 150, 156, 188, 189 and 194 (Noro and Nonaka 2014) - are shaded gold and pink, respectively (summarized in **Table 1**). Residues that do not match either lineage are shaded blue. Residue 31, which defines the PSMB8 type is shaded orange (A type with A or V), aqua (F type with F or Y), purple (S type with S or T) or green (K type). Sequence identifiers for ruddy bowfin and longnose gar are indicated with red and blue text— this is to highlight the novel PSMB8 S and K types observed in these holostean lineages. Sequence identifiers for two PSMB8 types from the divergent denticle herring (Denclu) are indicated with green text — note that, as both forms appear to have insertions or deletions between position 1 and 31 of the mature protein, position 31 of these sequences do not align with position 31 of all other species in this analysis making the assignment of a PSMB8 type challenging. Our nomenclature used denticle herring residues that align with position 31 of other species (F and S). In contrast, the actual residues in position 31 are K and D (green text). Sequences previously reported by Noro and Nonaka (2014) are indicated by asterisks (*). All sequences and display identifiers are provided in **Supplementary Tables S1 - S4**.

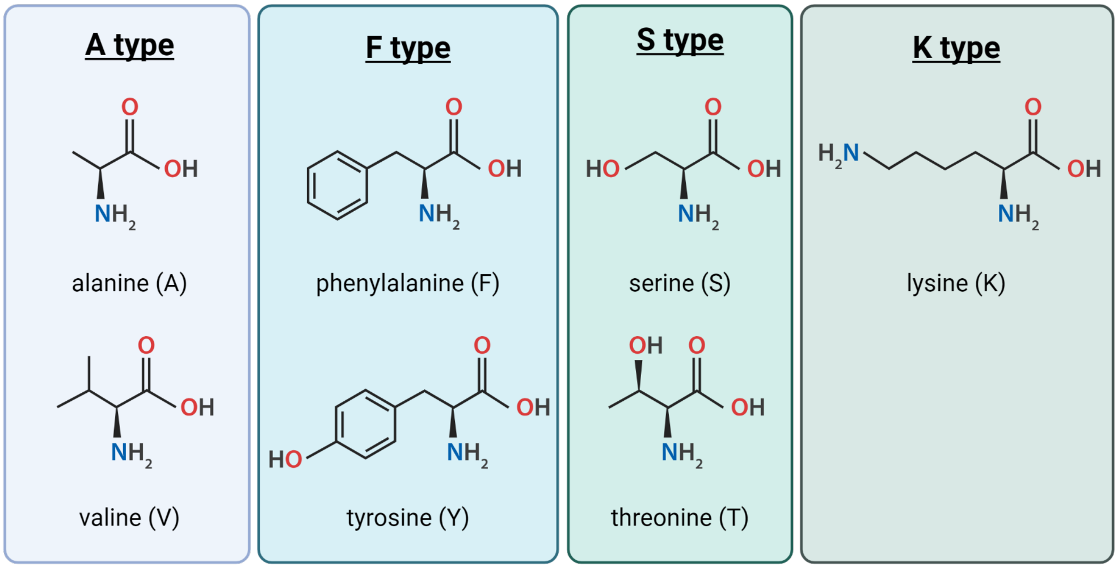

**Supplementary Figure S4. Thirty-first residues of PSMB8 types.**

Amino acids present in the 31st position of the mature PSMB8 protein define the "type". Figure generated using BioRender *(publication license and URL will be provided once accepted for publication)*.

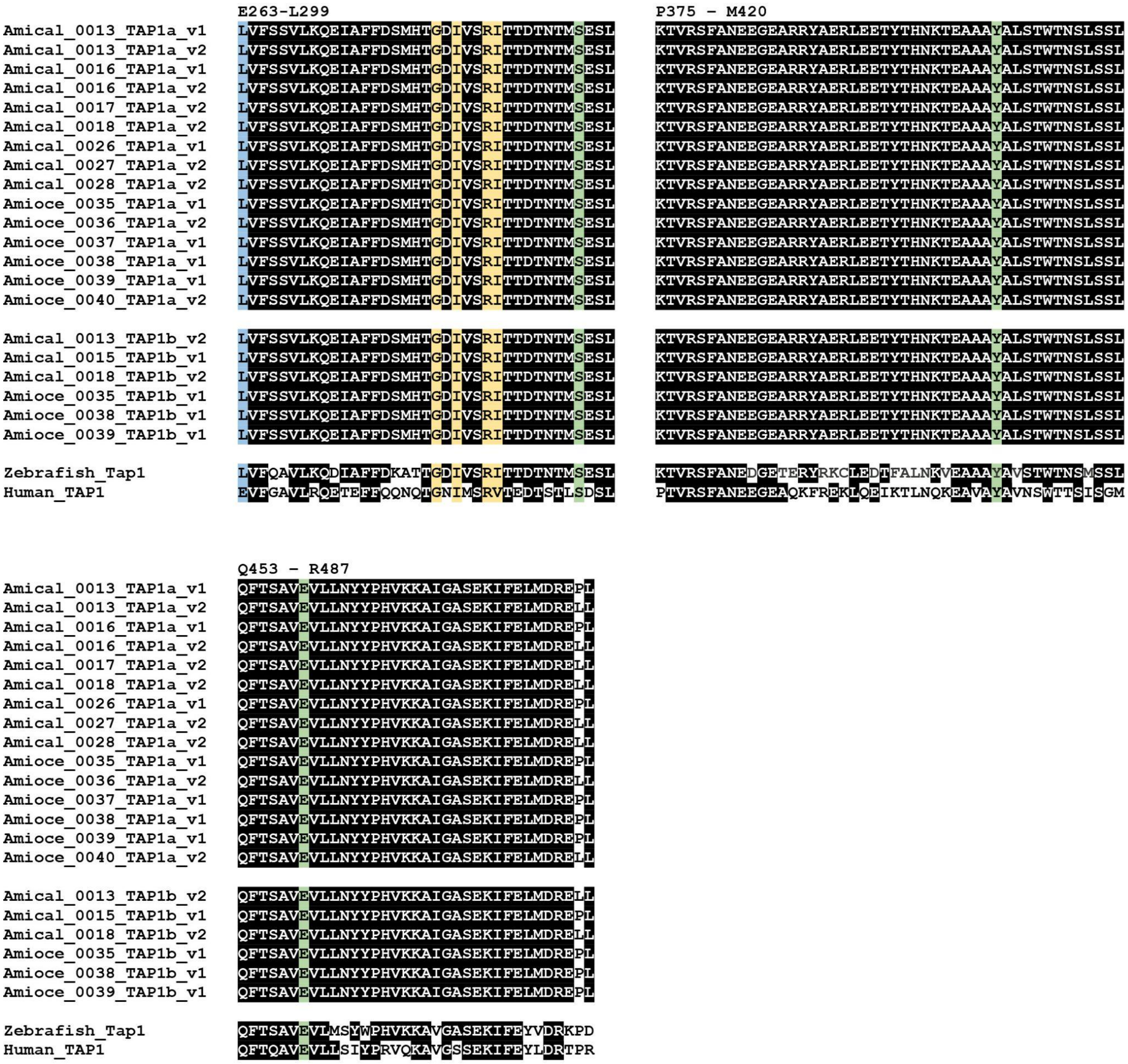

**Supplementary Figure S5. Essential TAP1 residues from individual bowfin.**

Bowfin TAP1 regions encoding residues related to functionality were aligned using Clustal Omega (Sievers and Higgins 2021). Human (GenBank NP_000584.3) and zebrafish (GenBank XP_002665053.1) TAP1 sequences are included for reference with amino acid numbering from human TAP1 shown above. Positions that are ≥90% identical are shaded black. Positions related to functionality (263), peptide sensing (282, 284, 287, 288), and substrate specificity (296, 408, 459) are shaded in blue, yellow, and green respectively (Lehnert and Tampé 2017).

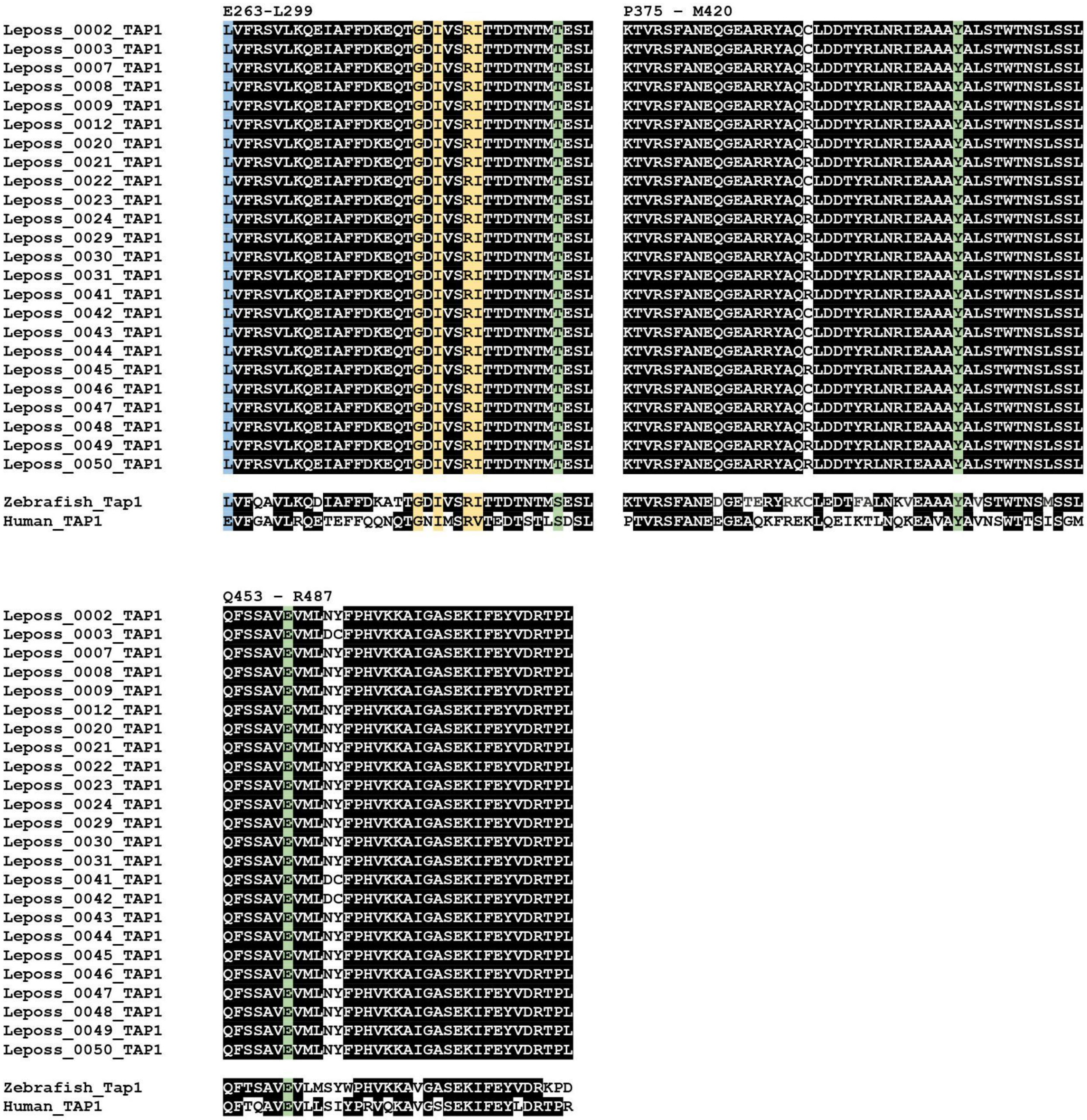

**Supplementary Figure S6. Essential TAP1 residues from individual longnose gar.**

Longnose gar TAP1 regions encoding residues related to functionality were aligned using Clustal Omega (Sievers and Higgins 2021). Human (GenBank NP_000584.3) and zebrafish (GenBank XP_002665053.1) TAP1 sequences are included for reference with amino acid numbering from human TAP1 shown above. Positions that are ≥90% identical are shaded black. Positions related to functionality (263), peptide sensing (282, 284, 287, 288), and substrate specificity (296, 408, 459) are shaded in blue, yellow, and green respectively (Lehnert and Tampé 2017).

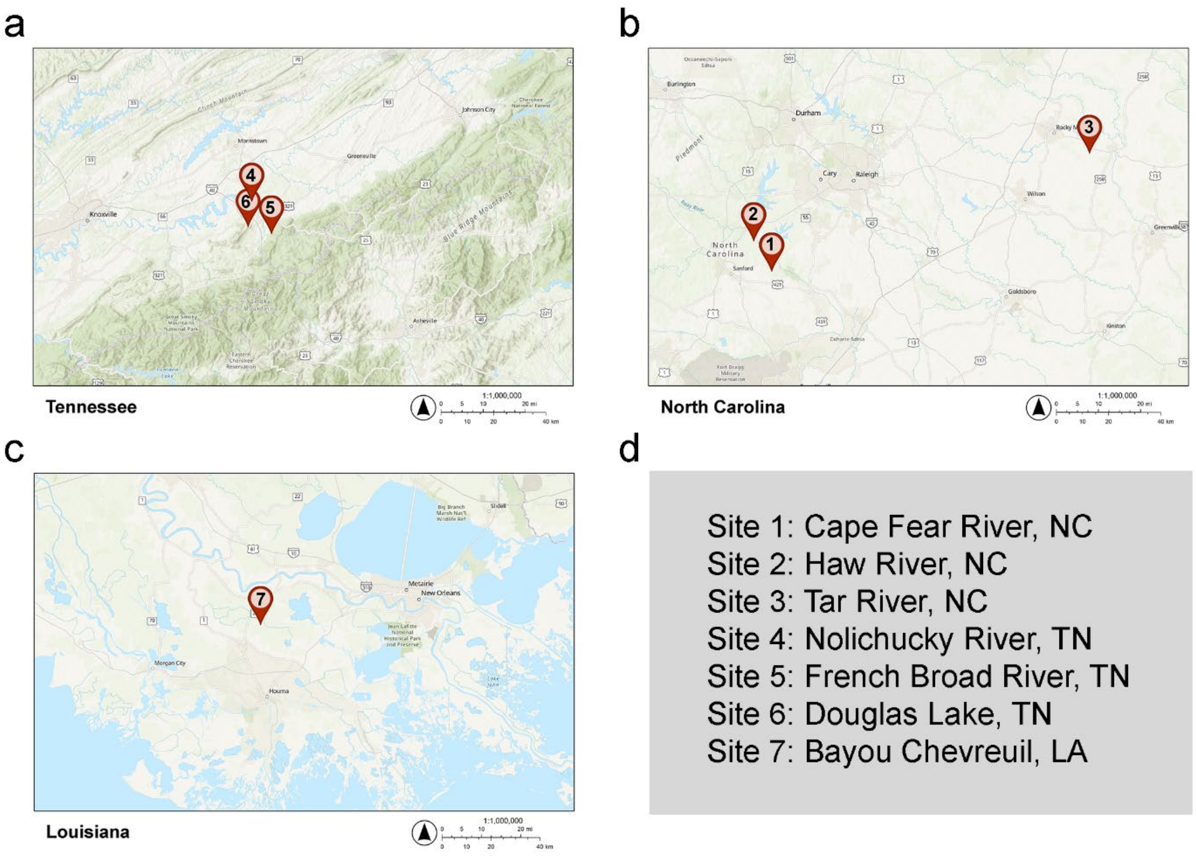

**Supplementary Figure S7. Collection sites for bowfin and longnose gar.**

The locations where bowfin and longnose gar were collected are indicated in maps of **a** Tennessee, **b** North Carolina, and **c** Louisiana. Locations are listed in **d** with coordinates provided in **Supplementary Tables S7 and S8**. Maps were generated using ArcGIS Online software.

**References.**

Bi X, Wang K, Yang L, Pan H, Jiang H, Wei Q, Fang M, Yu H, Zhu C, Cai Y, et al. 2021. Tracing the genetic footprints of vertebrate landing in non-teleost ray-finned fishes. *Cell* 184:1377–1391.e14.

Braasch I, Gehrke AR, Smith JJ, Kawasaki K, Manousaki T, Pasquier J, Amores A, Desvignes T, Batzel P, Catchen J, et al. 2016. The spotted gar genome illuminates vertebrate evolution and facilitates human-teleost comparisons. *Nat. Genet.* 48:427–437.

Lehnert E, Tampé R. 2017. Structure and dynamics of antigenic peptides in complex with TAP. *Front. Immunol.* 8:10.

Noro M, Nonaka M. 2014. Evolution of dimorphisms of the proteasome subunit beta type 8 gene (PSMB8) in basal ray-finned fish. *Immunogenetics* 66:325–334.

Sievers F, Higgins DG. 2021. The Clustal Omega Multiple Alignment Package. *Methods Mol. Biol.* 2231:3–16.
